# Extracellular electron transfer genes expressed by candidate flocking bacteria in cable bacteria sediment

**DOI:** 10.1101/2024.02.22.581617

**Authors:** Jamie JM Lustermans, Mantas Sereika, Laurine DW Burdorf, Mads Albertsen, Andreas Schramm, Ian PG Marshall

## Abstract

Cable bacteria, filamentous sulfide oxidizers that live in sediments, are at times associated with large flocks of swimming bacteria. It has been proposed that these flocks of bacteria transport electrons extracellularly to cable bacteria via an electron shuttle intermediate, but the identity and activity of these bacteria in freshwater sediment remains mostly uninvestigated. We coupled metagenomics and metatranscriptomics to 16S rRNA amplicon-based correlations with cable bacteria from two time series experiments up to 155 days. We identified bacteria expressing genes for extracellular electron transfer and motility, including synthesis genes for potential extracellular electron shuttles: phenazines and flavins. Of the 85 high quality MAGs (Metagenome Assembled Genomes >90% complete and <5% contaminated), 56 had genes encoding flagellar proteins, and of these 22 had genes encoding extracellular electron transport proteins. The candidate flockers constituted 21.4% of all MAGs and 42.1% of the proposed flocking bacteria expressed extracellular electron transfer genes. The proposed flockers belonged to a large variety of taxonomic groups: 18 genera spread across 9 phyla. Our data suggest that electric relationships in freshwater sediments between cable bacteria and other microbes likely help to generate and/or sustain cryptic element cycling and ‘deep oxygen breathing’, affecting more element cycles than sulfur, like metal– and in particular iron– and carbon cycles.

## Introduction

Cable bacteria are centimeter-long filamentous sulfide oxidizers that drastically change the environment when they are active (Rao et al, 2015; Pfeffer et al, 2012; Risgaard-Petersen et al, 2012). The cable bacteria respire oxygen in the oxic zone of sediments they live in, while oxidizing sulfide in sulfidic parts (Pfeffer et al, 2012). This results in physical separation of the oxidative and reductive metabolic processes, necessitating transport of electrons from sulfide oxidation through their filaments upwards to where the oxygen reduction takes place, making them an electric conduit (Meysman et al, 2019).

Cable bacteria belong to the genus *Candidatus* Electronema and *Candidatus* Electrothrix, which are part of the *Desulfobulbaceae* (Trojan et al, 2016). Thus far, cable bacteria have not been isolated in pure culture and remain in sediment enrichments or natural microbial communities (Thorup et al, 2021). In both natural settings and sediment enrichments, cable bacteria impact microbial community composition and activity. This has been observed in several instances such as chemoautotrophic sulfide-oxidizing *Gamma*– and *Epsilonproteobacteria* (/*Campylobacterota*) that accrued more carbon around cable bacteria, iron-reducers that appeared in higher abundance, or increased sulfate concentrations that stimulated sulfate reduction (Liau et al, 2022; Liu et al, 2021; Sandfeld et al, 2020; Otte et al, 2018; Vasquez-Cardenas et al, 2015). Some associations likely arose from the geochemical changes (Liu et al, 2021; Sandfeld et al, 2020). However, not all associations could be explained by these, which led to suggestions that electric interactions may occur to close metabolic cycles (Bjerg et al, 2023; Meysman, 2018; Vasquez-Cardenas et al, 2015).

An example of potential electric interactions with cable bacteria is the flocking behavior that occurs around *Ca.* Electronema aureum GS in freshwater sediment (Bjerg et al, 2023; Lustermans et al, 2023). In a microscopy slide, cable bacteria were discovered to have hundreds of bacteria swimming actively around individual filaments in a chemotactic manner, seemingly without touching the cable bacteria (Bjerg et al, 2023). Bjerg and colleagues suggested that flocking bacteria exhibit a form of extra-cellular electron transfer (EET) that may be electron shuttle mediated. Electron shuttles are compounds that can be oxidized and reduced without being consumed and can be used by microbes to respire insoluble electron acceptors such as metal particles or electrodes (Pham et al, 2008; von Canstein et al, 2008). The flockers would donate electrons to such a shuttle, followed by the cable bacteria coupling the oxidation of these shuttles to the reduction of O_2_ in the oxic zone several millimeters distant from the flocking (Bjerg et al, 2023). Through usage of EET, the flocks of bacteria would be able to survive in an anoxic environment without oxygen as terminal electron acceptor, but with possibly flavins, humic substances (abundant in many sediments) or other common shuttle substances in sediment facilitating electron transport between flocking bacteria and cable bacteria (Bjerg et al, 2023, Monteverde et al, 2018; Ratasuk and Nanny, 2007).

Knowledge about flocking is limited: it appears consistently with active, electron-conducting cable bacteria, the flockers are of diverse morphology and taxonomy, and flocking is suggested to be an electron-shuttle-based EET behavior, but the details of the EET mechanism remain unknown (Bjerg et al, 2023; Lustermans et al, 2023).

Several pathways used for EET are known from model organisms: *Shewanella oneidensis* MR-1, *Geobacter sulfurreducens*, *Sideroxydans lithotrophicus* ES-1, *Pseudomonas putida*, *Thermincola potens* JR, *Rhodopseudomonas palustris* TIE1, and *Enterococcus faecalis* (Paquete et al, 2022; Hederstedt et al, 2020; Pham et al, 2008). Ten porin-cytochrome complexes (pccs) (2 Mtr’s, Pio, Mto, Cyc2, TherJR_2595, OmabcB and its three homologues) that export electrons and oxidize substances outside of the cell have been described (Paquete et al, 2022; Shi et al, 2016). EET appears widespread, making it likely that other microorganisms have different pathways to perform EET (Baker et al, 2022; Zhong and Shi, 2018; Shi et al, 2016; 2014). Utilizing shuttle-based EET requires the presence of shuttles; if not already in the environment, shuttles could be excreted by microorganisms (Monteverde et al, 2018; Glasser et al, 2017; Newman and Kolter, 2000). Even nanomolar concentrations of shuttles present in the sediment can explain the observed flocking phenomenon (Bjerg et al, 2023). For shuttle synthesis and excretion, specialized proteins are used such as PhzABCDEF, IpdG and MexGHI-OpmD for phenazines, and Bfe, YeeO and RibBA/X for flavins (Kotloski and Gralnick, 2013; McAnulty and Wood, 2014; Blankenfeldt and Parsons, 2015; Sakhtah et al, 2016; Brutinel et al, 2013).

With this knowledge, a microbial community’s potential for EET may be determined through presence and expression of EET-encoding genes using metagenomics and metatranscriptomics. We combined this with previously and newly published 16S rRNA amplicon data collected over 76-155 days (Lustermans et al, 2023) to determine whether the microbial community in a cable bacteria enrichment has EET potential. We determined specific taxonomic groups that correlate with *Ca.* Electronema and looked at their potentials and usage of EET.

## Materials and methods

The samples originate from 3 time series experiments with 2 different sediments: 2 separate time series of 79 (TS1) and 155 days (TS2) using autoclaved lake sediment (Vennelyst Park, Denmark) inoculated with an enrichment of *Candidatus* Electronema aureum GS (Thorup et al, 2021) and a 79-day time series with sieved Brabrand Lake sediment. The GS enrichment has been maintained as a laboratory enrichment (Thorup et al, 2021), the Brabrand lake sediment was collected specifically for this study.

16S rRNA sequences derived from the time series with *Ca.* Electronema aureum GS were used from Lustermans et al. (2023) with exception of the 155-day time point, which was added, but collected and generated similarly as described by Lustermans et al. (2023).

### Sediment collection and microsensor set-up

Black (sulfidic) freshwater sediment was retrieved from Brabrand Lake, Denmark (56.140458, 10.144352) at a water depth of 3-4 m. The sediment was stored with overlying water at 15 °C for 2 weeks to ensure anoxia and remove fauna.

Before incubation, sediment was homogenized, sieved (pore size: 0.5 mm) and distributed into core liners that were closed with a rubber stopper at the bottom. Sedimentation occurred after 24 h, after which the sediment surface was aligned with the core liner edge and cores were then submerged in autoclaved tap water. The aquarium was kept at 15 ᵒC, covered with aluminum foil to prevent algae formation, equipped with air circulators and a lid to prevent excessive evaporation. Overlying water was replenished and refreshed multiple times during incubation.

Microsensor measurements for O_2_ and EP (electric potential), to determination the active current-generating (sulfide-oxidation) zone were performed and analyzed as described in Lustermans et al. (2023).

### 16S rRNA amplicon community analysis

The taxonomically classified collection of V3-V4 16S rRNA amplicons were generated after total RNA extraction in a previous study (Lustermans et al, 2023) or produced in this study using the same procedure. Correlation analysis was carried out on i) the complete set (TS1, TS2) of *Ca.* Electronema aureum GS samples, ii) all natural samples (Brabrand Lake, *Ca.* Electronema sp.), and iii) subsets of the cable bacterial growth phase to avoid spurious correlations resulting from increased relative abundance of other 16S rRNA sequences relative to cable bacteria. Growth was defined as increase of cable bacterial abundance: T3-T33 for the first set of amplicons (TS1) and T2-T27 for amplicons from the second time series (TS2) (Lustermans et al, 2023). Fractional abundances of ASVs (amplicon sequence variants) were calculated, followed by pruning of ASVs unclassified to genus level. The remaining classified ASVs were then agglomerated into genera with Phyloseq v1.42.0 (McMurdie & Holmes, 2013). Spearman correlations were performed at genus-level against *Ca*. Electronema (sp.), followed by a p-value adjustment to account for multiple testing (R v4.2.2; cor.test (method = spearman), p.adjust). Heatmaps were generated using ampvis2 v2.6.4 (Andersen et al, 2018).

### Metagenomics

Two metagenomes derived from the *Ca.* Electronema aureum GS enrichment were used: a short-read Illumina dataset from a trench-slide incubation where flocking was observed (Bjerg et al, 2023), and a long-read Oxford Nanopore metagenome from anoxic bulk sediment (Sereika et al, 2023). A trench slide consists of two glued-together microscopy slides, with a central cavity in the upper one. This is filled with sediment, covered with anoxic tap water and a cover slip (Bjerg et al, 2023). Cable bacteria glide into an observable plane towards inwards-diffusing oxygen. Sulfide diffuses from the cavity.

Preprocessing, sequencing, and binning of the metagenomes was carried out as described previously (Bjerg et al, 2023; Sereika et al, 2023), with the resultant metagenome-assembled genomes (MAGs) incorporated into this study.

FastANI v1.32 (Jain et al, 2018) was run against MAGs from both sources to remove duplicates. If two near-identical (>98.5% ANI) MAGS were found, the most complete MAG was chosen for further analysis as calculated by CheckM2 v1.0.1 (Chklovski et al, 2022). Genes of each MAG were identified and annotated using PROKKA (Seemann, 2014), followed by taxonomic classification of the bins with GTDB-tk v2.1.0 and database r207 (Chaumeil et al, 2019). The final bin-collection was run through KOfamscan v1.3.0 (Aramaki et al, 2019) with a 1E-6 cut-off value and compared with Blastp (Camacho et al, 2009) with a 1E-6 cut-off value to an inhouse prepared database containing EET proteins from various origins (Table S1).

### Metatranscriptomics

Metatranscriptomes were prepared of three timepoints (TS1: days 3, 26, 33; Lustermans et al, 2023). Samples represent low and high cable bacteria activity/abundance for comparison. Total RNA was extracted in duplicate from the active current-generating zone (anoxic bulk sediment) using the RNeasy PowerSoil Total RNA kit (Qiagen) and concentrated with the kit RNA Clean & Concentrator (Zymo Research, USA) including an in-column DNase treatment (Ambion, USA). Metatranscriptome sequencing was done (2x 50 bp, 50 mio reads/sample) with Illumina HiSeq (Illumina, USA) without prior rRNA removal (DNASense, Denmark). The sequences were trimmed, and size filtered using Trimmomatic v.0.39 (Bolger et al, 2014).

### Mapping and *de novo* analyses

Trimmed and filtered sequences were mapped against the final MAGs with bbmap v39.01 (Bushnell 2010). This was then used for differential gene expression analysis with DESeq2 v1.39.8 (R v4.2.2) (Love et al, 2014). The expressed genes were compared to the previously mentioned Blastp analysis of the metagenomes and results were compiled (Table S1-3).

*De novo* assembly was done on the pooled metatranscriptome sequences using rna-SPAdes v3.15.4 (Bushmanova et al, 2019). Trimmed sequences were mapped against the assembly with bbmap v39.01 to determine transcript abundance (Bushnell, 2010). Protein-coding genes were identified in assembled transcripts using FragGeneScan v1.30 (Rho et al, 2010). Genes were annotated and identified as EET-genes as described for the metagenomes. Taxonomy of genes was determined with eggNOG-mapper v2.0.1 (Cantalapiedra et al, 2021). Discovered EET-genes were compared to the metagenome and correlating taxa.

## Results

### 16S rRNA amplicon analyses

We acquired 36 samples from the *Ca.* Electronema aureum GS time series. These originated from two time series: TS1 (76 days, 9 time points) and TS2 (155 days, 10 time points). From the Brabrand Lake sediment we generated 12 samples over 76 days (9 time points). All samples had more than 3909 reads with an upper limit of 132888 reads and a median of 22202.

Positive correlations were determined between rRNA of the genus *Ca.* Electronema and other genera in TS1 and TS2 and in the natural sediment from Brabrand lake, which contained several strains of *Ca.* Electronema.

In the inoculated enrichment *Ca.* Electronema aureum GS rRNA increased in relative abundance to ∼55% of the community on days 26-33, after which it decreased slowly to day 155 without disappearing (Fig 1). The genera *Desulfosporosinus*, *Dehalobacter*, *Papillibacter*, *Acetanaerobacterium*, and *Desulfoprunum* positively correlated throughout the enrichment. *Christensenellaceae* R-7 group, *Desulfovibrio*, *Papillibacter*, *Anaerovorax*, and *Ruminiclostridium* correlated positively during the cable bacteria’s growth phase (day 3-33).

**Figure 1.**
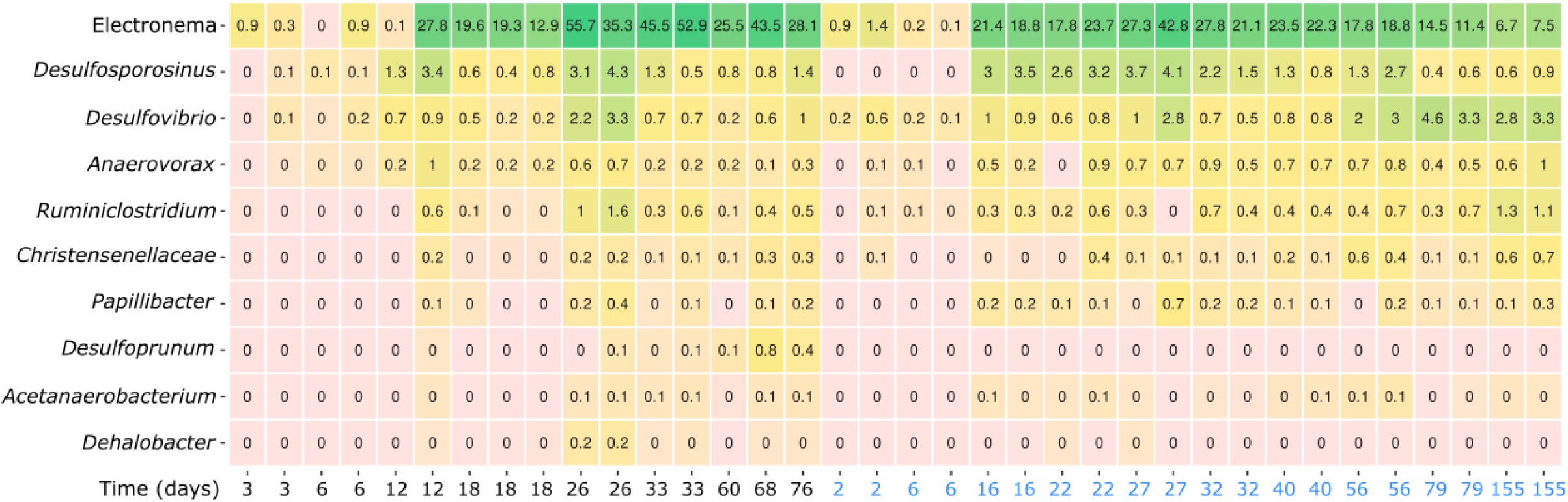
– Heatmap of both time series containing *Ca.* Electronema aureum GS enriched sediment showing the genera that correlated positively with *Ca.* Electronema aureum GS over time (2-155 days, black TS1, blue TS2). Numbers on heatmap show percentage abundances.

Different correlations were found when the two time series were separated into TS1 and TS2 (SI Fig 1 & 2). In TS1; *Dehalobacter*, *Ruminiclostridium*, *Christensenellaceae* R-7 group, *Desulfurispora* and *Acetanaerobacterium* correlated positively with cable bacteria during the 76 days of incubation. In TS2, *Desulfosporosinus* was the sole positive correlation with cable bacteria over 155 days.

In the natural sediment from Brabrand Lake, the cable bacteria, *Ca.* Electronema sp., rRNA showed lower relative abundance than in the GS enrichment with a maximum of 1.1% on day 64 (Fig 2). The positive correlations that were found in the natural cable bacteria cores during 76 days were with: *Desulfonema*, *Sulfurimonas*, *Desulfatirhabdium*, CL500-29 marine group *(Ilumatobacteraceae*), and *Cuspidothrix* LMEYCA_163.

**Figure 2.**
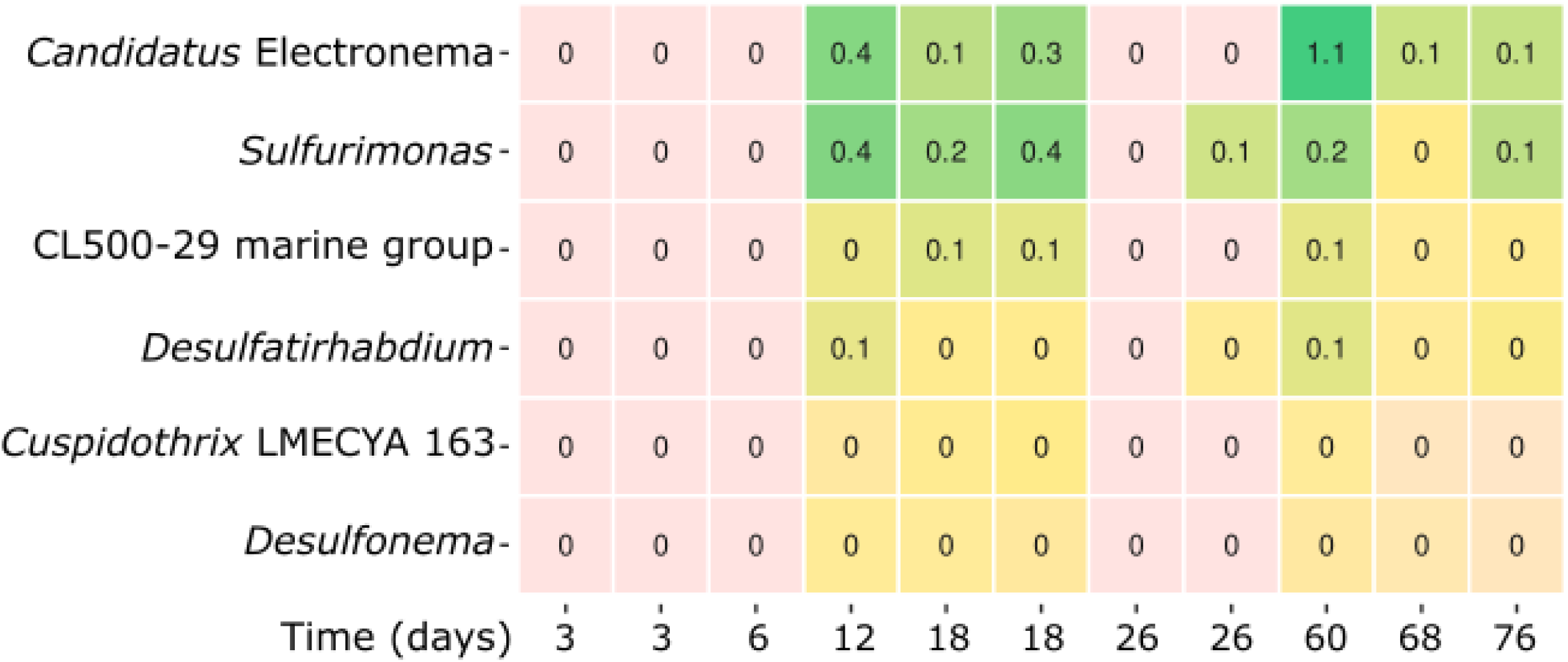
– Heatmap of naturally occurring *Ca.* Electronema sp. in Brabrand Lake, showing the genus-level taxa that correlated positively with cable bacterial succession over time (3-76 days). Numbers on heatmap show percentage abundances.

### Metagenomics and –transcriptomics analyses

We determined presence or absence of genes for motility, EET or electron-shuttle related genes in the metagenomes derived from the *Ca.* Electronema aureum GS enrichment and observed whether transcripts of these genes could be detected by RNA sequencing. The metagenomes were binned into 103 unique MAGs. We identified 5083 EET-, shuttle synthesis– and excretion genes in the MAGs, to which 959 genes were mapped from the metatranscriptomes (Table S1). Only 164 of these overlapped with the *de novo*-assembled metatranscriptomes, which yielded another 534 unmapped EET-related genes. To determine whether EET-related transcripts were more prevalent when there were more cable bacteria (days 26, 33) compared to when they were in lower abundance (day 3), significant differences in transcript abundance of outer membrane EET– or electron shuttle genes were analysed for both the MAG-mapped and the *de novo*-assembled transcripts (Fig 3). As expected, transcripts for 79% of *Ca.* Electronema aureum GS genes were significantly more abundant when the bacterium was in high abundance (Table S2). Gene percentages of MAGs that showed significantly more abundant transcripts during high cable bacterial abundance (days 26, 33) versus low abundance (day 3) varied widely: 41% – <0.5% (Table S2). There were 22 out of the 103 MAGs that had no significantly more abundant transcripts compared between the cable bacteria abundances (Table S2).

**Figure 3.**
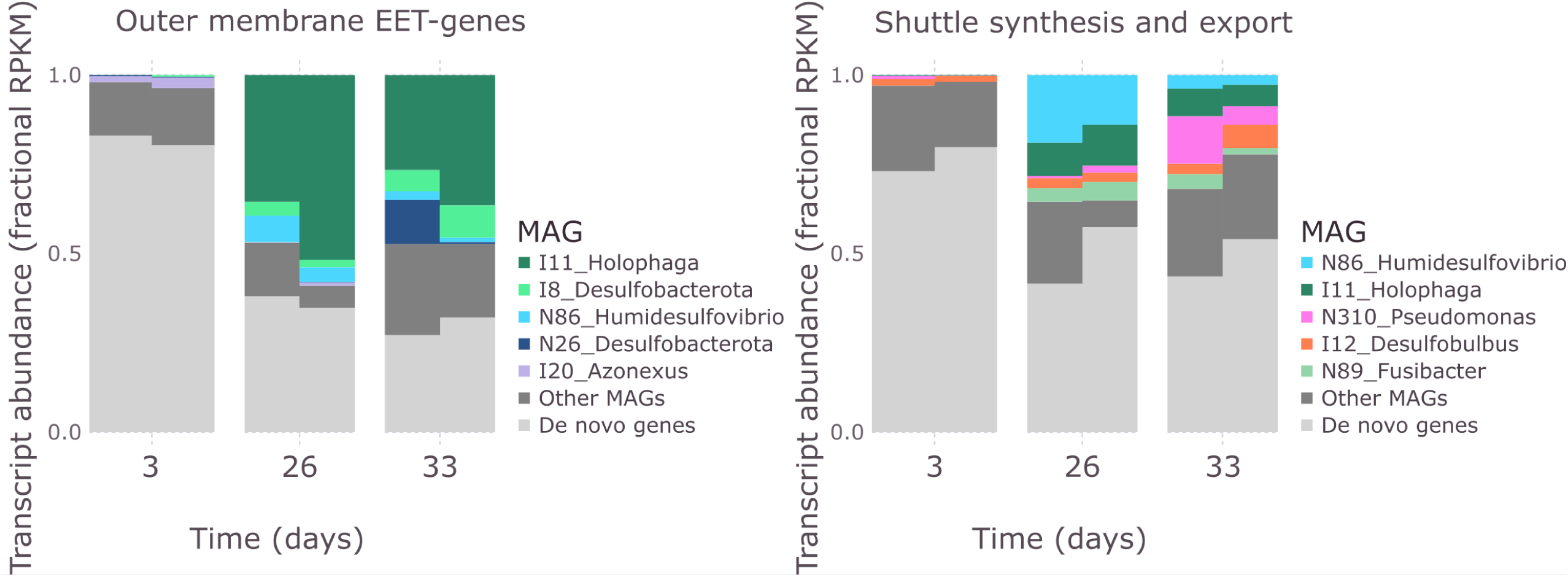
– Fractional abundances of porin cytochrome complexes (pccs)– and electron shuttle-genes transcribed during low (day 3) or high (days 26,33) relative cable bacterial abundance. Only the five highest scoring MAGs were shown individually, other are clustered in ‘Other MAGs’, ‘De novo transcripts’ comprises genes that were not mapped to any MAG. The pcc genes included: *ndh3, eetB, dmkA, omabcB, omabcC, therJR_2595, therJR_0333, therJR_1122, cyc2, mtrABC, mtrDEF, pioAB, mtoAB, dmsAEF, cwcA, extEFG, extBCD, omcA, omcS, omcZ*. And the shuttle genes used were: *yeeO, bfe, ribBA, ribBX, ribD, ribE1, ribE2, ipdG, phzDEFG, mexGHI, OpmD*.

### Porin cytochrome complexes (pccs) and outer membrane cytochromes (omcs)

Complete sets of pcc encoding genes, that are necessary for microbes to perform EET, were found in 18 bacterial genomes (Fig 4). A single bin contained four copies of the *cwcA* gene that spans the cell-wall in the Gram-positive EET-performing bacteria *Thermincola* sp. Incomplete versions of pccs were found in many MAGs (Table S1). In total we found 22 potential EET-performing bacteria. Pccs (including *dmkA*, *ndh3, eetB* and *cwcA, cyc2* as 3 complete pccs, excluding *mtoD* and *pioC* as these are not part of the membrane-bound part of the porin complex) were transcribed by 57 MAGs of the 103 total MAGs (Table S1). 54 of these 57 MAGs transcribed at least one gene of a pcc on either day 3, 26 or 33. However, many transcribed pcc or omc-genes that were detected in the *de novo* assembly did not match to any MAG (Fig 3). I11_Holophaga transcribed the highest amount of outer membrane EET-genes (pccs and omcs), but only during high cable bacteria abundance (days 26, 33; Fig 3). I8_Desulfobacterota, N86_Humidesulfovibrio, N26_Desulfobacterota and I20_Azonexus transcribed outer membrane EET-genes as the others in the top 5 MAGs and were found to do so both at low and high cable bacterial abundance (Fig 3). Most outer membrane EET-genes that were identified only in the *de novo* assembled metatranscriptomes were seen on day 3, when cable bacterial abundance was low (Fig 3).

**Figure 4.**
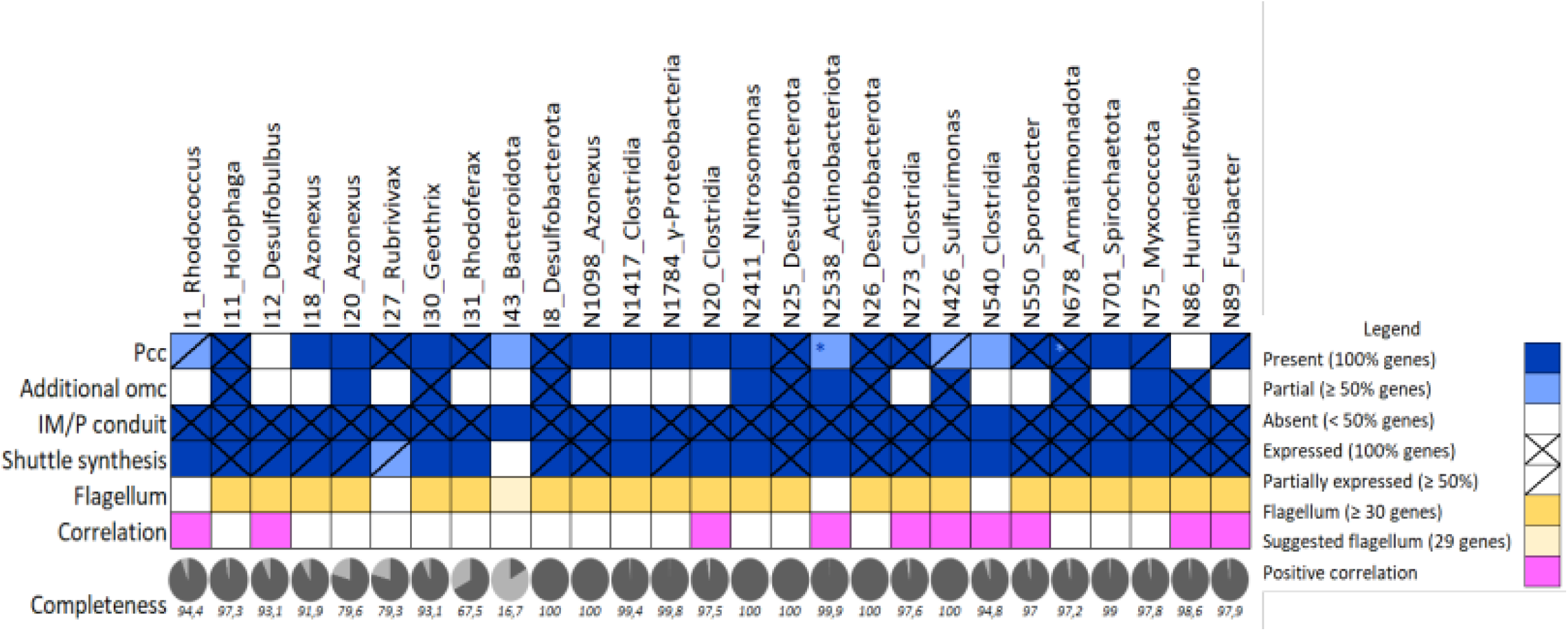
– Selection of MAGs identified as potentially flocking around cable bacteria due to EET-capabilities or positive correlation with cable bacteria. Presence/absence of motility genes and positive 16S rRNA-based correlations (correlations were based on 16S rRNA genus level, rather than with the individual MAG). Completeness of the genomes, below, in percentages. “Additional omcs” (outer membrane cytochromes) include any gene that is an omc (*pioA, cyc2, mtoA, cwcA, omcA,B,C,S,Z, therJR_2595, extC,D,F, mtrA,F*).

Genes encoding cytochromes on the outside of the outer membrane (*omcA, omcS, omcZ*, and *therJR_2595*), were present in 22 MAGs and transcribed by 6 bacteria (Fig 4, Table S1). *OmcA* was found in only one MAG, all others were identified in at least two different MAGs.

*Desulfobacterota_*I8 transcribed *therJR_2595, omcS* and *omcZ* (Table S1). Over all, omcs were identified in 31 MAGs and transcripts of one or more omc were mapped to 15 MAGs (Table S1).

All MAGs classified as *Clostridia* (a genus positively correlated with cable bacteria according to 16S rRNA data) had the ability to express *eetB* and two or more other *E. faecalis*-type pcc-genes (Fig 4, Table S1). The *Sulfurimonas* expressed most genes found in the *S. oneidensis* pathway, *pioA*, and *mtoA* which are all parts of the pccs that appear essential for EET (Fig 4, Table S1). Significantly transcribed EET genes (pccs, omcs, P/IM cytochromes) during high cable abundance, belonged to 7 MAGs (I11, I18, I20, I8, N1098, N273, N89).

### Periplasmic (P) and inner membrane (IM) cytochromes

Aside from a pcc, microbes need a conduit to transport electrons from the cytoplasm to the outer membrane. Most MAGs encoded and, with few exceptions, transcribed *macA, cbcL, imcH, menB, menE,* and *fccA/3* genes for this purpose (Fig 4, Table S1). While other conduits: *cymA, imdcA, pdcA, imoA* were only sporadically transcribed and/or encoded (Table S1). IM/P conduits were transcribed by 78 out the 103 MAGs (Table S1).

### Electron shuttles and motility

Electron shuttle secretion and synthesis genes were transcribed by 64 MAGs (80 including the MexGHI-OpmD pump, which is not exclusively used for excretion of electron shuttles). Flavin production based on the *ribBA/X* gene was transcribed by 27 bacteria and 1 archaeon. Exporters for flavins encoded by *bfe* or *yeeO* were transcribed by 12 different bacteria (Table S1). One of the phenazine synthesis genes, *phzE*, was transcribed by 25 bacteria, while 54 bacteria transcribed the *ipdG* phenazine regulator gene of *P. aeruginosa* (Table S1). Both archaeal bins, methanogens, were able to produce electron shuttles, of which one transcribed two genes for shuttle production (Table S1). Six of the positively correlated bacteria transcribed synthesis and excretion genes for flavins or phenazines (Fig 4). The cable bacteria transcribed genes for flavin– and phenazine production (Fig 4). Electron shuttle synthesis gene expression (including *mexGHI, OpmD*), additional to those mapped to the MAGs, was also discovered in the *de novo* assembly (Fig 3). N86_Humidesulfovibrio transcribed the highest amount of shuttle synthesis-related genes, after what was found in the *de novo* assembly. Transcribed shuttle synthesis genes from I11_Holophaga, N310_Pseudomonas, I12_Desulfobulbus were identified at low and high cable bacterial abundance (Fig 3).

N89_Fusibacter, the 5^th^ highest shuttle synthesis/export-gene transcribing MAG, only transcribed these during high cable bacterial abundance (days 26, 33; Fig 3). As for the outer membrane EET-gene transcripts, higher percentages of electron shuttle synthesis/export-genes transcripts were identified in the *de novo* assembly during day 3, compared to days 26, 33 (Fig 3).

Out of the 103 MAGs, 66 bacteria were likely motile by flagella since they contained more than 30 of the 55 needed genes according to the KEGG database (Table S1).

## Discussion

We discuss specific taxonomic groups that were found to correlate with *Ca.* Electronema aureum GS and looked at their potential and expression for EET and electron shuttle synthesis. As we expect that flocking bacteria are a subset of these taxa, we determined whether motile bacteria could perform EET based on their genetic potential and expression. A candidate flocker needs: i) motility, ii) EET genes and transcription thereof, and iii) we consider 16S rRNA or transcript-correlations with cable bacteria.

For ii) we focused on complete pcc-homologs to a model EET-organism (Paquete et al, 2022; Hederstedt et al, 2020; Light et al, 2018; Shi et al, 2016; Carlson et al, 2011; Castelle et al, 2008). Not only pccs of model organisms may be functional since Conley et al. (2020) showed that *Vibrio natriegens* uses a hybrid EET pathway combining proteins from *Aeromonas* and *Shewanella*. We assumed that discovered hybrid pccs were functional if a complete outer-membrane-crossing complex could be created.

Flocking bacteria, exclusively found around actively electron conducting cable bacteria, were proposed to use electron shuttles and subsequently cable bacteria as alternative electron acceptors and calculated that concentrations of 2-187 nM need to be present (Bjerg et al. 2023). Thus small volumes of electron shuttles could sustain flocking, making presence of synthesis and excretion genes in the community expected. Many of the MAGs expressed flavin-synthesis gene *ribBA/X* combined with *yeeO* or *bfe* for export (McAnulty and Wood, 2014; Brutinel et al, 2013; Kotloski and Gralnick, 2013). Phenazine production was less common, but phenazine gradients remain plausible since many bacteria expressed *phzE* and *ipdG* (Glasser et al, 2017; Blankenfeldt & Parsons, 2014; Pham et al, 2008). The sediment origin, a eutrophic pond with plants, inhabited by animals, suggests that humic substances, a naturally occurring quinone-group-containing electron shuttle was present (Glasser et al, 2017; Klüpfel et al, 2014; Pham et al, 2008; Ratasuk and Nanny, 2007; Kappler et al, 2004).

We identified 22 candidate flockers, 21.4% of all MAGs. The anticipated active flocking community was 42.1% of all identified candidate flockers. Candidate flockers were taxonomically classified in 9 phyla and 18 different genera. This large variety of taxonomies fits with the observed morphological diversity (Bjerg et al, 2023; Lustermans et al, 2023). Many of these genera have previously been observed in sediment enrichments of cable bacteria or associated with them (Sachs et al, 2022; Liu et al, 2021; Liau et al, 2020; Sandfeld et al, 2020; Otte et al, 2018; Vasquez-Cardenas et al, 2015).

From the many different interesting taxa with regards to EET and shuttle synthesis, we discuss the top four phyla of interest with regards to EET capabilities and correlation with cable bacteria. Other taxa (divided per phylum) can be found in the supplementary information (Note S1).

### Acidobacteriota

Two *Acidobacteriota*; a *Holophaga*, and a *Geothrix* were identified as motile. The *Holophaga* expressed the *extEFG* pcc from the model organism *Geobacter*, the omc *mtrF* (*Shewanella*) and inner conduits that *Geobacter* and *Shewanella* – two well-described Fe-respiring bacteria-have (Table S1; Shi et al, 2016). *ExtEFG* was significantly expressed (p-values ranged from 3.83E^-08^ to 0.00276) during high cable bacterial abundance (days 26 and 33), highlighting the importance of this pcc for the *Holophaga*. It also transcribed the highest amount of outer membrane EET-genes from all MAGs, especially much during high cable bacteria abundance (days 26, 33), which suggests that I11_*Holophaga* associates on an electrical basis with the presence of cable bacteria.

The *Geothrix* showed the same potential but expressed only *mtrF* and *fccA*, a conduit. However, absence of observed transcripts is not equal to absence of the protein. *Holophaga* and *Geothrix* have been implicated in EET before and seen in bioanode communities (Olmsted et al, 2022; Arbour et al, 2020). The *Holophaga* and *Geothrix* may well perform EET as flockers. This may be an interesting synergy where they consume the ferrous iron from FeS dissolution (Risgaard-Petersen et al, 2012), stimulated by cable bacteria, and then donate the gained electrons to cable bacteria, previously also suggested by Otte et al (2018).

### Campylobacterota

Only 1 *Campylobacterota* was identified amongst the MAGs. Colleagues observed that chemoautotrophic sulfur-oxidizing *Epsilonproteobacteria* correlated with– and were active in absence of oxygen around cable bacteria (Liau et al, 2022; Liu et al, 2021; Otte et al, 2018; Vasquez-Cardenas et al, 2015). Direct association with cable bacteria has thus far not been found. The *Campylobacterota*, a *Sulfurimonas*, had pcc-, shuttle-synthesis– and conduit genes. No pcc was complete, but with genes for *pioA, mtoA, dmsA* and *dmsE*, it may be using an undetected porin for transmembrane electron-transport. Based on 16S rRNA amplicons, the *Sulfurimonas*-genus correlated positively with the cable bacteria in the natural lake sediment, suggesting an association. A recently discovered *Sulfurimonas* was shown to oxidize sulfide by transferring electrons to manganese(IV)-oxide particles, which requires a form of EET due to the solid nature of this electron-acceptor (Henkel et al, 2019). The incomplete pccs of the *Sulfurimonas* leave its potential for IET up for question. De novo assembled EET genes were classified as *Campylobacterota*. As we found only one MAG in our two individually and differently generated metagenomes, we assumed that these belonged to the *Sulfurimonas*. The EET-genes were mostly transcribed when cable bacteria were highly abundant, suggesting that it was interacting with cable bacteria or used EET during changes brought on by cable bacterial activity.

### Desulfobacterota

Five motile *Desulfobacterota* expressed inner conduits, four expressed omcs, and three potential flockers (I8, N25, N26) had genes for 2-4 pccs as were found in *G. sulfurreducens*, of which 2-3 were expressed (Table S1). These three were *Pseudopelobacteraceae*, the family that *Geobacter* belongs to and which thus contains well-studied EET performers (Table S5; Rotaru et al, 2014; Smith et al, 2014; Shresta et al, 2013; Butler et al, 2010). I8 significantly (p.adjust = 0.00499) expressed *omcZ* on days 26 and/or 33 when relative abundance of cable bacteria was high and was the second highest outer membrane EET-gene-transcribing MAG (Fig 3). Thus it is likely intensively using these proteins during high cable bacterial abundance. As described above, cable bacteria have a complex interaction with the iron cycle, providing several different ways that they could stimulate the activity of an iron-oxidizing or –reducing microorganism (Otte et al, 2018). Cable bacteria activity acidifies the sediment, releasing Fe(II) from FeS (Risgaard-Petersen et al, 2012). Fe(II)-oxidation could be coupled to the reduction of the cable bacteria, which would then produce Fe(III) for conventional Fe(III)-reduction. Or one could imagine that iron-cycling microorganisms are not involved in iron-cycling at all, but rather couple the oxidation of organic matter to the reduction of cable bacteria.

Additionally, cable bacteria have also been shown to indirectly stimulate sulfate-reducing bacteria in freshwater sediments (Liu et al, 2021; Sandfeld et al, 2020). That indirect relationship could potentially be visible in the correlation analyses as positive correlations with the cable bacteria. Two *Desulfobacterota* genera were found to positively correlate with cable bacteria in the enrichment and two in the natural sediment, this would confirm the previously shown correlations. However, such correlations do not exclude *Desulfobacterota* from having EET-genes nor from being a potential flocking bacterium. Sulfate reduction does not necessarily exclude the potential for IET as some species can use organics, fumarate, AQDS (two electron shuttles), iron, electrodes, and potentially the cable bacteria as alternative electron acceptors (Pecheritsyna et al, 2012; Cordas et al, 2008).

I12_*Desulfobulbus* and N86_*Humidesulfovibrio*, were sulfate-reducers that correlated with cable bacteria (16S rRNA) and that showed up in the top 5 MAGs that transcribed outer membrane EET-genes or shuttle synthesis genes (Fig 1-3). Neither contained potential for a pcc, but N86 expressed an omc (*therJR_2595*) and *dmsA*. Both expressed synthesis genes for flavins and phenazines. The *Humidesulfovibrio* significantly expressed (p.adjust = 0.0029-0.0353) three *dmsA* and two *ipdG* genes (p.adjust = 0.0082-0.0084) during high cable bacterial abundance (days 26, 33), suggesting that these were used for an important process. It may be able to extra-cellularly reduce DMSO, like *Shewanella* (Gralnick et al, 2006). This is, however, unlikely to occur in the suboxic zone. We suggest that *Humidesulfovibrio* may help produce shuttles but not perform IET as we expect of the flockers (Bjerg et al, 2023). Our data strengthens earlier observations where *Desulfobacterota*-groups could be interacting with cable bacteria in the anoxic zone (Liau et al, 2022; Liu et al, 2021; Sandfeld et al, 2020; Vasquez-Cardenas et al, 2015). *De novo* assembled EET– and shuttle-synthesis gene transcripts during high cable bacterial abundance were classified as *Desulfovibrionales* and *Desulfobacterales*, that may be used to support IET. Unfortunately these were not mapped to any MAGs, which shows that even with two metagenomes, the resolution is not high enough to discover all the relevant EET-related bacteria. The *Desulfovibrionales* and *Desulfobacterales* orders have been associated with cable bacteria before, where they were linked based on indirect interactions around sulfate reduction (Liu et al, 2021; Sandfeld et al, 2020). Thus, there might be direct cooperation between cable cells and these candidate flockers in the suboxic zone where there is very little sulfide available (Pfeffer et al, 2012; Risgaard-Petersen et al, 2015). Sulfate is limited in freshwater environments, yet with the influx from cable-bacterial-sulfide oxidation, many sulfate reducers will be found here and competition for sulfate likely arises (Liu et al, 2021; Sandfeld et al, 2020). Alternatively, these *Desulfobacterota* may not be reducing sulfate at all, but rather directly using the cable bacteria as an electron acceptor. These candidate flockers may use IET as a way to avoid competition with other sulfate reducers (Pecheritsyna et al, 2012; Cordas et al, 2008).

### Firmicutes

A wide range of *Firmicutes* have been observed in bio-electrochemical systems, observed individually as electroactive and studied as model organisms (Arbour et al, 2020; Hederstedt et al, 2020; Paquete, 2020; Ishii et al, 2018; Light et al, 2018). *Firmicutes* were also previously seen enriched in freshwater cable bacteria sediments (Sachs et al, 2022; Liu et al, 2021). We observed that they are enriched in our sediments, and many were even correlating with cable bacteria growth in the enrichment and the natural sediment, confirming that the previously found enrichments of *Firmicutes* are likely based on a form of interaction. Most of the amplicon-based positively correlating taxa belonged to the *Clostridia*. Amongst the eight *Clostridia* MAGs, most EET-genes were transcribed on day 3, during low cable bacterial abundance. The genes belonged almost exclusively to the Gram-positive pcc (*dmkA*, *eetB*, *ndh3*) or shuttle synthesis (Hederstedt et al, 2020).

N273_*Clostridia* expressed its pcc completely and N89_*Fusibacter* expressed *dmkA* and *ndh3*. These, and N550_*Sporobacter* also transcribed flavin and phenazine-synthesis genes. *Fusibacter* belonged to the top 5 MAGs that expressed the most shuttle-synthesis genes, which it did mostly during high cable bacterial abundance. N89_*Fusibacter* likely used flavins as intermediaries for IET as those were the expressed genes. It may fulfill an important role by sustaining electron shuttle concentrations.

Based on the presence of a pcc we indicate *Sporobacter* (N550) and *Fusibacter* (N89) and four *Firmicutes* of the *Clostridia* or *Bacillus* classes (N1417, N20, N273, N1017) as candidates flockers.

### Gammaproteobacteria

We identified 28 *Gammaproteobacteria* of which 18 could perform chemoautotrophic sulfur-oxidation, based on their genome (Bjerg et al, 2023). Previously, chemoautotrophic sulfur-oxidizing *Gammaproteobacteria* have been observed correlating with– and being active in absence of oxygen around cable bacteria (Liau et al, 2022; Liu et al, 2021; Otte et al, 2018; Vasquez-Cardenas et al, 2015), but a direct association with cable bacteria has not yet been shown. The limited access to high quality electron acceptors (oxygen and nitrate/nitrite) where their donors are means that these bacteria needed an alternative electron acceptor (Meysman, 2018). As it is expected that sulfide concentrations are low around cable bacteria, these bacteria would actively have to move towards reduced sulfur sources, or may take advantage of the increased sulfate reduction rates (Sandfeld et al, 2020). Most of the *Gammaproteobacteria* MAGs had genes for shuttle synthesis and some expressed them of which N310_*Pseudomonas*, was the third shuttle-synthesis-gene transcriber, suggesting that it was actively producing riboflavin and phenazine, especially on day 33 (Fig 3, Table S1). Four MAGs had a complete homolog of *E. faecalis’* pcc (I18_*Azonexus*, I9_*Azonexus*, N1098_*Azonexus*, N1784_*Gammaproteobacteria*). Two MAGs contained *mtoBAD* genes like *S. lithotrophicus* (I20_*Azonexus,* I31_*Rhodoferax*). The *Sideroxydans* MAG (I37) did not contain this pcc, it only contained (and expressed) *mtoD* and was thus not included as a candidate flocker. MAG I27_*Rubrivivax* expressed *dmsA, mtrBA* and *mtrF* which may form a functional hybrid complex where *mtrF* would replace *mtrC* as omc (Conley et al, 2020). The reduction window of *mtrC* (–500-+100mV) is wider than *mtrF’s* (–400-+50mV, Paquete et al, 2022), thus dependent on which shuttles were used, reduction of the shuttles could occur (Paquete et al, 2014). Another potential hybrid complex was identified in N2411, a motile *Nitrosomonas* with *mtoA, therJR_2595,* and *dmsF*, where *dmsF* would function as the integral OM-protein, anchoring *mtoA* from the inside, with *therJR_2595* as omc. Under anoxic conditions, *Nitrosomonas europaea* and *N. eutropha* use organics such as acetate, lactate or pyruvate coupled to nitrite as electron acceptors (Schmidt, 2009). We suppose that N2411_*Nitrosomonas* can use these organics too, but coupled to reducing the cable bacteria as there is no nitrite available in the suboxic zone (Marzocchi et al, 2020). We suppose that these hybrid pccs could be functional and include *Nitrosomonas* and *Rubrivivax* as possible flockers. The potential flockers are four *Azonexus* species (I18, I9, N1098, I20), *Nitrosomonas* (N2411), *Rhodoferax* (I31), and *Rubrivivax* (I27). Three of these MAGs (I20, I27, I31), expressed their pcc genes and most transcribed inner conduits. The *Rubrivivax* did not contain enough flagellar assembly genes to assume flagellar motility (23/55), however this MAG lacked ∼20% of its genome and the type strain (*R. gelatinosus*) was described to be motile (Willems et al, 1991).

Therefore, we assumed that it was capable of motility. The genera we found as potential flockers have members which have been verified as EET-performers: *Sideroxydans, Azonexus, Rhodoferax* (Jain et al 2022; Jangir et al, 2016; Finneran et al, 2003).

### Conclusion and ecological impact

We assumed that flockers would correlate with cable bacteria, due to their metabolic interaction (Lustermans et al, 2023). Our data reinforces this, but also showed that exceptions could exist. In addition there is a large volume of EET-gene transcripts found in the metatranscriptomes, but not mapped to any MAG, suggesting that the diversity of potential flockers is likely larger than the 22 we propose here. The large amount of genes and the expression thereof, combined with the low required concentration for electron shuttles, support the hypothesis that flockers use shuttles as intermediates to transport electrons to cable bacteria (Bjerg et al, 2023; Meysman, 2018). Based on taxonomy, the flockers are an extraordinary metabolically flexible group ranging from organotrophs to iron-(metal-) and sulfur-users, but electron-donor origins need experimental confirmation. The overlap in metabolic functions of genera that positively correlated with cable bacteria between our sediment enrichment and the natural lake sediment shows that the simplified community in the enrichment still represents natural processes, albeit in potentially lower abundances. The discovery of a large group of likely EET-performing bacteria that could associate with cable bacteria, extends the influence of oxygen reduction deep into anoxic sediments. While cable bacteria are only known to directly oxidize sulfide, a direct electric association with other microorganisms widens the range of possible electron donors that could be oxidized in anoxic cable-bacteria-containing sediment to include metals and organic compounds. This means that the impact of cable bacteria on carbon and metal cycling could be much greater than previously thought.

## Acknowledgements

We thank Susanne Nielsen, Britta Poulsen, Lykke Bamdali, Marie Rosenstand Hansen and Marie Braad Lund for excellent technical laboratory assistance and troubleshooting with amplicon sequence analysis. We would like to thank Casper Thorup for fruitful scientific discussions.

This research was financially supported by the Danish National Research Foundation (DNRF136), the Flanders Research FWO (S004523N), the University of Antwerp via the TopBOF program, and the Poul Due Jensen Foundation (Microflora Danica).

## Data availability

16S rRNA amplicon sequences from all three time series experiments are available at NCBI in the Sequence Read Archive (SRA) and can be found under accession number PRJNA837365 and PRJNA1058976. Metagenomic data is available at the NCBI database under accession number PRJNA730231 (Bjerg et al, 2023) and from ENA under bio project number PRJEB52550 (Sereika et al, 2022). Metatranscriptomic data is available from the NCBI SRA-database under accession number PRJNA1057540.

## Author contributions

JJML, LDWB, AS and IPGM conceived the study

JJML, IPGM, LDWB, MA, AS and MS designed experiments

JJML, MS and LDWB performed experiments

JJML, IPGM and MS analyzed data

JJML wrote the manuscript with input from all authors

